# PlotsOfData – a web app for visualizing data together with its summaries

**DOI:** 10.1101/426767

**Authors:** Marten Postma, Joachim Goedhart

## Abstract

Reporting of the actual data in graphs and plots increases transparency and enables independent evaluation. On the other hand, data summaries are often used in graphs since they aid interpretation. State-of-the art data visualizations can be made with the ggplot2 package, which uses the ideas of a ‘grammar of graphics’ to generate a graphic from multiple layers of data. However, ggplot2 requires coding skills and an understanding of the tidy data structure. To democratize state-of-the-art data visualization of raw data with a selection of statistical summaries, a web app was written using R/shiny that uses the ggplot2 package for generating plots. A multilayered approach together with adjustable transparency offers a unique flexibility, enabling users can to choose how to display the data and which of the data summaries to add. Four data summaries are provided, mean, median, boxplot, violinplot, to accommodate several types of data distributions. In addition, 95% confidence intervals can be added for visual inferences. By adjusting the transparency of the layers, the visualization of the raw data together with the summary can be tuned for optimal presentation and interpretation. The app is dubbed PlotsOfData and is available at: https://huygens.science.uva.nl/PlotsOfData/

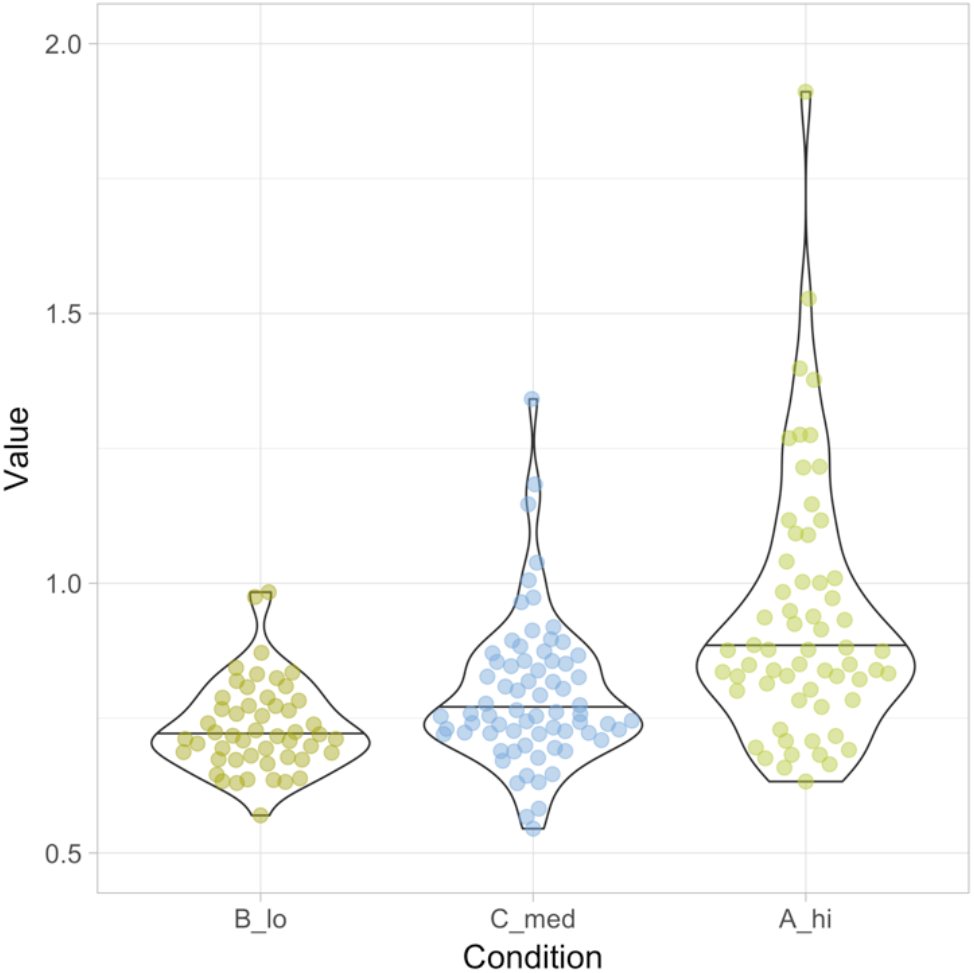

## Introduction

Over the recent years, several groups have advocated the presentation of the actual data in graphs instead of data summaries (Weissgerber et al., 2015; Rousselet et al., 2017; Drummond and Vowler, 2011). Raw data can be visualized in different ways, including histograms and dotplots. Data summaries may be displayed to aid interpretation of the data. In addition, direct comparison of the different categories/conditions can be done by ‘visual inference’ if 95% confidence intervals are supplied (Cumming and Finch, 2005; Cumming et al., 2007; Gardner and Altman, 1986).

Several commercial software packages are available to draw data and their summaries. However, ideally, such tools should be open source, freely available and allow contributions or modifications by users. One example of a free open source web-based app to plot a combination of raw data and summaries is BoxPlotR (http://shiny.chemgrid.org/boxplotr/). The web based app is described in a paper (Spitzer et al., 2014) that is remarkably well-cited. Its popularity reflects a demand for easy-to-use applications that generate publication quality data visualizations. However, this popular online tool is skewed towards boxplots as data summaries and has hardly any options for customizing the combined display of data and summaries. Moreover, the plots are rather basic in appearance.

State-of-the art data visualization is possible with the R package ggplot2 that uses the ideas of a ‘grammar of graphics’ to generate a graphic by using multiple layers of data (Wickham, 2011). The multilayered approach enables to compose a graph from individual components, each of which can be independently adjusted. The option to apply transparency to the data layers adds to the flexibility. Yet, the high-quality data visualization provided by ggplot2 requires coding skills and understanding the concept of tidy data (Wickham, 2014).

To democratize state-of-the-art data visualization of raw data with a selection of statistical summaries, we generated a web tool that we dubbed PlotsOfData. The web tool uses ggplot2 to compose the graphs and handles data in ordinary spreadsheet (wide) format as well as the tidy data format. Since creating graphs with PlotsOfData does not require coding skills, the high-quality data visualization provided by ggplot2 is now available to anyone. Some of the features of PlotsOfData will be highlighted below.

## Availability & code

PlotsOfData is available at: https://huygens.science.uva.nl/PlotsOfData/

The app uses the shiny package and was written in R using R (https://www.r-project.org) and Rstudio (https://www.rstudio.com). It uses several freely available packages (shiny, ggplot2, dplyr, tidyr, readr, magrittr, ggbeeswarm, readxl). The source code is attached as a supplement and an up-to-date version is available at Github together with information on how to install and run the app locally: https://github.com/JoachimGoedhart/PlotsOfData

## Data input and structure

The data can be provided by copy paste into a text box or by upload of two file formats, i.e. csv (comma separated values) or xls (excel) format. Several delimiters (comma, tab, semicolon, space) are recognized. Two example files are available in the app for testing the application. These files are also available as csv files (https://github.com/JoachimGoedhart/PlotsOfData). The native structure that ggplot2 uses is the ‘tidy’ format (Wickham, 2014) and this data structure is accepted. However, data is often stored in a wide, spreadsheet-type structure in which each column reflects a condition. This wide format is the default data structure that is used by the app. Users may select columns from the spreadsheet data that should not be included in the graph. After the input, the wide data is converted into tidy format, assuming that each column is a condition with a single row header that lists each condition. To promote the understanding of tidy data, the data can be downloaded in a tidy format.

## Data visibility

The data is shown as transparent dots. Offset can be added to the dots to avoids overlap for larger number of dots. When quasirandom is selected as the offset, the dots are shown according to the data distribution (i.e. similar to a violinplot). Both the offset and user-defined visibility of the raw data can be adjusted to optimize the visualization of the raw data. For low numbers of data, it is pertinent to plot the data (Weissgerber et al., 2015; Drummond and Vowler, 2011; Vaux, 2014). For very large numbers of data, i.e. when the dots show substantial overlap, one may consider to make the data fully transparent and only plot its distribution with a violinplot.

## Statistical summary

Any of four statistics can be added to summarize the data, i.e. median, mean, boxplot or violinplot. The median is not sensitive to outliers and as such a robust indicator of the central value (Wilcox and Rousselet, 2017, 2018). The median is also indicated in the boxplot (Krzywinski and Altman, 2014; McGill et al., 1978) and violinplot by a horizontal line. Since both boxplots (Krzywinski and Altman, 2014) and violinplots (Hintze and Nelson, 1998) reflect data distribution, they are only appropriate if sufficient data is provided (the lower limit is now set at n=10).

To enable inference by eye, the 95% confidence interval (CI) can be added to the plot (Cumming and Finch, 2005; Cumming et al., 2007; Gardner and Altman, 1986). For boxplots, the 95% CI is indicated by notches (Mcgill et al., 1978; Krzywinski and Altman, 2014). The original definition of notches was reported by McGill et al., but its calculation does not correct for small sample size (McGill et al., 1978). Therefore, notched boxplots should be used with care for smaller samples (n<20).

The 95% CI that is calculated when the median or violinplot is selected is calculated by bootstrapping (1000 samples) and determining the 95% CI from the 2.5th and 97.5th percentile (Wood, 2004). Since bootstrapping requires a representative sample from the population, it is only suitable if sufficient data is present. Since the underlying population that was sampled from is unknown, it is per definition, unclear what ‘sufficient’ means. To reduce the chances that the 95% CI does not correctly reflect the population we have set the minimum number of data points in the app at 10 per condition for the calculation of the 95% CI.

## Transparent layers

The raw data can be combined with any of four data summaries, i.e. mean, median, boxplot, violinplot. In addition, the 95% CI can be added for inferences. The simultaneous visualization of the data and statistics achieved by using (transparent) layers. For optimal visualization the order of the layers is defined as follows (from first to last, last appearing on top): 1) box- or violinplot, 2) raw data, 3) mean or median, 4) 95% CI. The visualization can be optimized by user defined transparency of the layers.

## Table with Statistics

The statistics that are selected for visualizing the data are also documented in a table on a separate tab. The default statistics that are listed in the table are depending on the summary statistics that are shown in the graph. For instance, when the mean is selected the mean and standard deviation are shown in the table but when the median is selected the median and the median absolute deviation (MAD) are included. The user can change the default statistics that are shown in the table and re-arrange their order by drag-and-drop of the columns. Moreover, the number of digits that are shown can be adjusted. The table can be exported to pdf or copied to the clipboard. Table 1 is an example output of the summary related to the data that is shown in figure 1.

**Figure 1.**
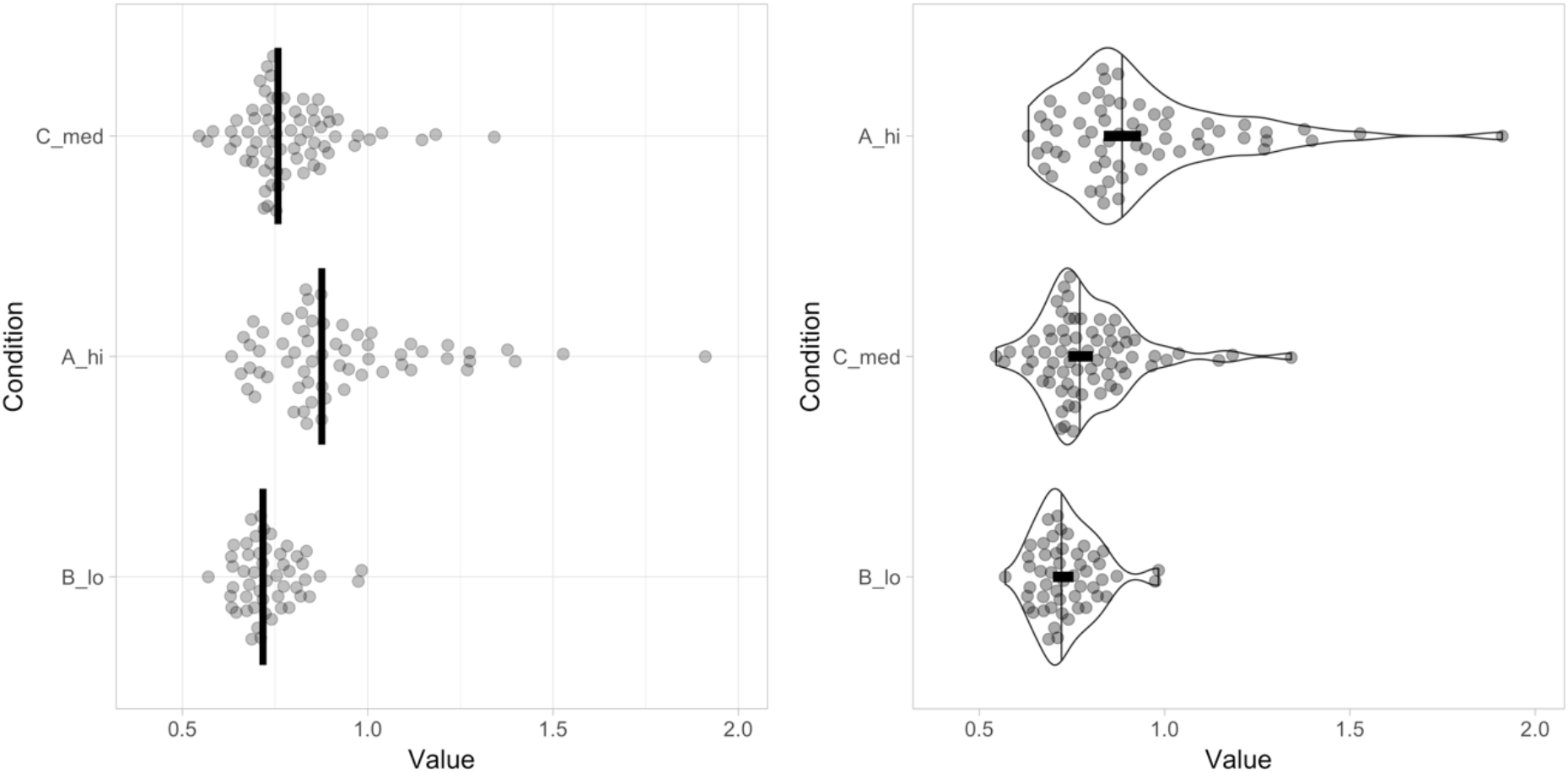
Visualization of the sample data provided by the app, showing the data and summary from three conditions. The left panel shows the data as dots and the median with a vertical line. The right panel shows the data (sorted according to median values) as dots, the distribution with a violinplot and the 95% confidence of the median by a black horizontal bar. Both graphs are rotated to improve readability of the conditions.

**Table 1:**
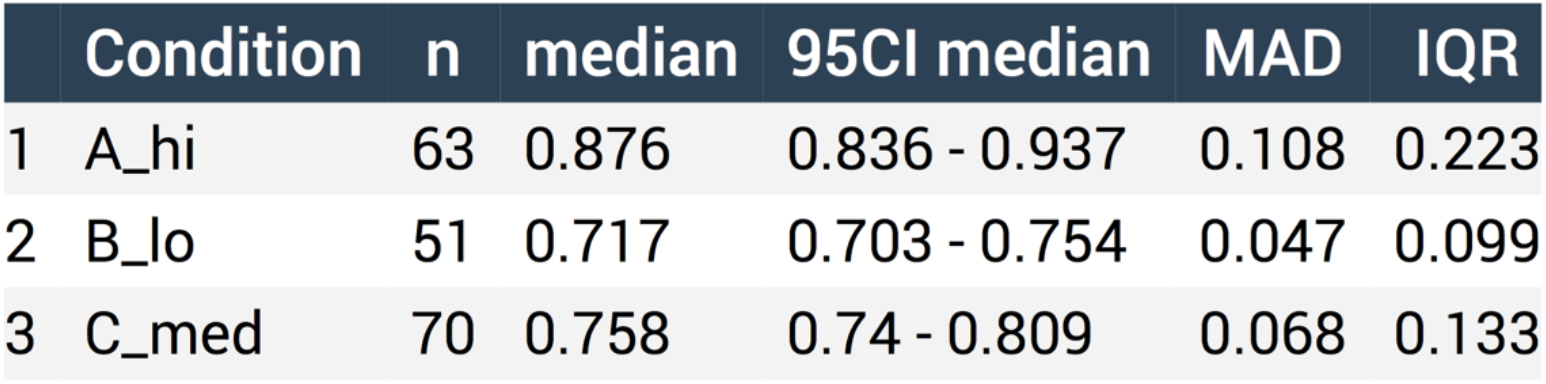
Summary of statistics related to the data shown in figure 1

## Color

The default graph is in black and white. Color can be added to either the data, the statistics or both. There is an option to use the standard palette, or several colorblind safe palettes that are optimized for categorical, qualitative data (https://personal.sron.nl/∼pault/). Finally, user defined colors can be added by using color names or hexadecimal codes.

## Data sorting

The categories can be sorted in three different ways. First, the data can be visualized in alphabetical order, which is the default order for ggplot2. Second, the categories can be shown in the same order as provided (by copy paste or in the uploaded file). Third, the data can be sorted according to the median value.

## Output

The graphs can be directly saved from the web browser that is running the app (e.g. by drag and drop from the web browser). Alternatively, two options are available for downloading the figure, pdf and png. The pdf can be edited by software that handles vector-based graphics for further adjustment of the lay-out.

## Conclusion

The shiny app was made with the aim to enable anyone to visualize their data in combination with a selection of summaries. The user defined mixing of dotplots with statistical summaries should improve the creation of graphs and visual inferences. The use of boxplots, violinplots and 95% CI requires sufficient data. It is however not agreed upon what ‘sufficient’ implies (Vaux, 2014; Weissgerber et al., 2015; Wilcox and Rousselet, 2017). In the app, the minimum is set at n=10 for showing boxplots, violinplots and 95%CI, but it is up to the user to critically assess whether this is sufficient. The source code for the app is available and the threshold can be readily changed in the code. Regardless of the statistics that is shown, it is recommended to plot the data for low to medium n (Weissgerber et al., 2015; Drummond and Vowler, 2011; Wilcox and Rousselet, 2017; Vaux, 2014).

Finally, we hope that the high-quality plots created with PlotsOfData will improve transparent communication of scientific data which will be beneficial for both researchers and their audience.

## Acknowledgments

PlotsOfData is inspired by BoxPlotR (http://shiny.chemgrid.org/boxplotr/). The code for the shiny app is partially derived from ggplotGUI (https://github.com/gertstulp/ggplotgui/) by Gert Stulp. The colorblind safe palettes were developed by Paul Tol (https://personal.sron.nl/∼pault/). We are grateful to Auke Folkerts (UvA, The Netherlands) for help with the server that runs shiny.

